# Rare earth elements extraction from Idaho-sourced surface soil by phytomining

**DOI:** 10.1101/2024.08.05.606409

**Authors:** Kathryn Richardson, Amin Mirkouei, Kasia Duellman, Anthony Aylward, David Zirker, Eliezer Schwarz, Ying Sun

**Author notes:** Corresponding authors: E-mail addresses (K. Richardson) and (A. Mirkouei)., Mail address: Tingey Administration Building, Suite 312, University of Idaho, Idaho Falls, 83402, USA.

## Abstract

Environmentally-friendly and low emission extraction methods are needed to meet worldwide rare earth element (REE) demand. Within a greenhouse setting, we assessed the REE hyperaccumulation ability of four plant species (e.g., *Phalaris arundinacea, Solanum nigrum, Phytolacca americana*, and *Brassica juncea*) and the impact of amending REE-rich soil with biochar or fertilizer and watering with citric acid solution. Harvested samples were pyrolyzed, and the resulting bio-ores were acid-digested and underwent elemental analysis to determine REE content. Amending soil with fertilizer and biochar increased bio-ore production, while plant species explained most variation in bioaccumulation factor. *Phalaris arundinacea* achieved the highest average REE concentration of 27,940 ppm for targeted REEs (i.e., cerium, lanthanum, neodymium, praseodymium, and yttrium) and 37,844 ppm for total REEs. We successfully extracted REE-rich bio-ore from plant biomass and determined that soil amendment and plant species will be critical parameters in design and implementation of Idaho-based REE phytomining operations.

**Graphical Abstract:** 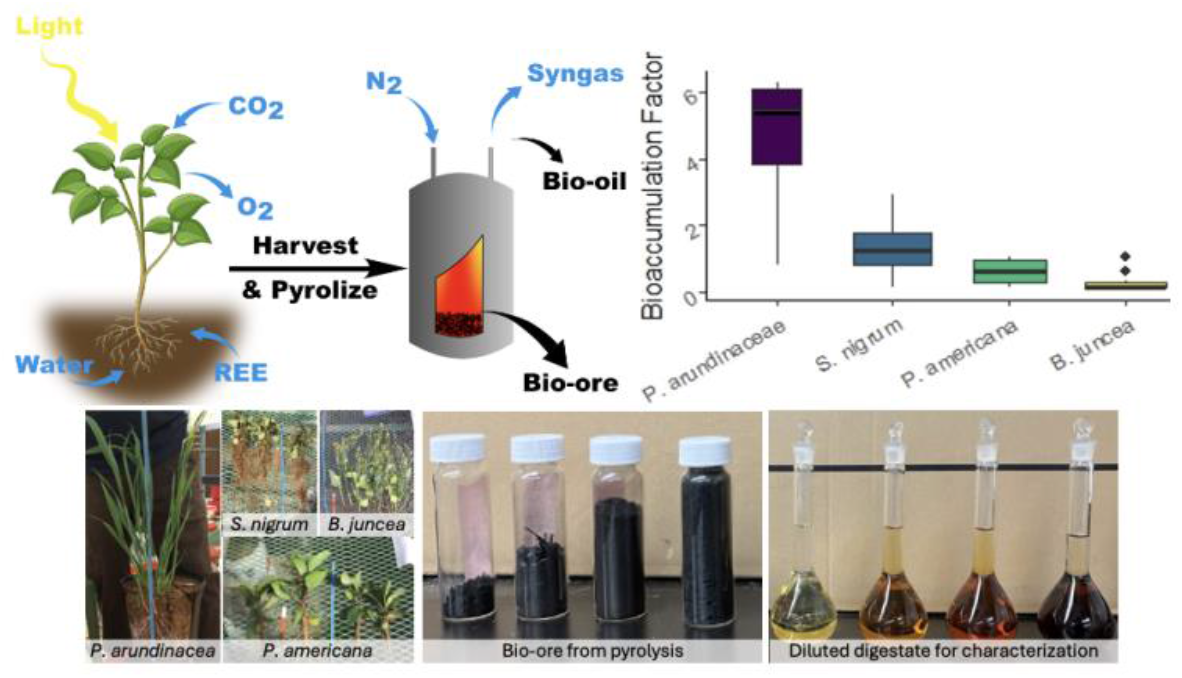

## Main

Across the globe there is a growing demand for products and technologies that rely on the use of rare earth elements (REEs). China and the United States of America (USA) are at the forefront of REE-based technology development, with demand for the elements spanning across medical devices, aircraft components, and communication systems, as well as renewable energy, rechargeable batteries, and smartphones ^1–3^. Mining of REEs within the USA is largely limited to a single California site, and most of the country’s supply is dependent on negotiations with China, which currently supplies the USA with most of the REEs it uses ^4,5^. The mining process also results in undesired consequences such as environmental degradation and the production of toxic mining waste that is often not processed further ^6^.

Idaho (a state in the northwestern USA) is endowed with a high abundance of REEs dispersed within soil that has not yet been targeted by any commercial mining operations ^1,7^. REEs comprise over 1% (11,601 ppm) of the total soil composition, especially across central Idaho in Lemhi Pass and Diamond Creek before crossing the Idaho-Montana state line (Supplementary **Table S1**). For comparison, soil from northern China contains only 0.024% (241 ppm) REEs and regolith originating in the southeastern USA is 0.105% (1,048 ppm) REEs ^8,9^. Idaho REE-rich soil can be an ideal candidate for phytomining, an alternative mining process used to recover REEs for processing and commercial use. Large scale phytomining operations in this region can address national needs for rare earth metals (REMs) production, along with job creation and other economic benefits for the region and nation. In addition, phytomining is a relatively cheap solution that can be integrated with traditional mining and improve soil quality for future crop production, forestation, and seeding of metal-intolerant native species to increase native biodiversity and avoid erosion ^10–12^.

REEs can enter vascular plants by forming complexes with organic macromolecules that facilitate adsorption to the root surface via transport proteins. They are transported via both symplastic and apoplastic pathways to the xylem, where they travel up the plant to the aboveground tissues via the transpiration stream and are then accumulated intracellularly, associating with multiple organelles and systems ^13^. Hyperaccumulator plant species tolerate high concentrations of REEs and other metals in their cells without suffering from metal toxicity ^13^. Bioaccumulation factor (BF) represents the concentration of target elements (e.g., REEs) in dried plant tissue versus the growth substrate (i.e., soil), with higher BF values indicating greater hyperaccumulation ability ^14^. A plant is typically considered a hyperaccumulator when it displays a BF value greater than one ^10^. These plants may be grown specifically for their REE storage abilities and then harvested and processed via pyrolysis (heating in the absence of oxygen) to convert plant biomass into bio-ore, rich in concentrated minerals, such as REEs ^10^. The 91-99% of minerals retained in bio-ore after pyrolysis can then be extracted and refined for commercial use via organic acid leaching (bioleaching) and metallurgy ^10,15–17^.

Due to the wide variety of plant species, substrates, growing treatments, and targeted elements, it is not fully understood which combinations of these factors yield the best phytomining results. To help motivate and direct the design of our experiments, we reviewed past phytomining research (**Table 1)**.

**Table 1.**
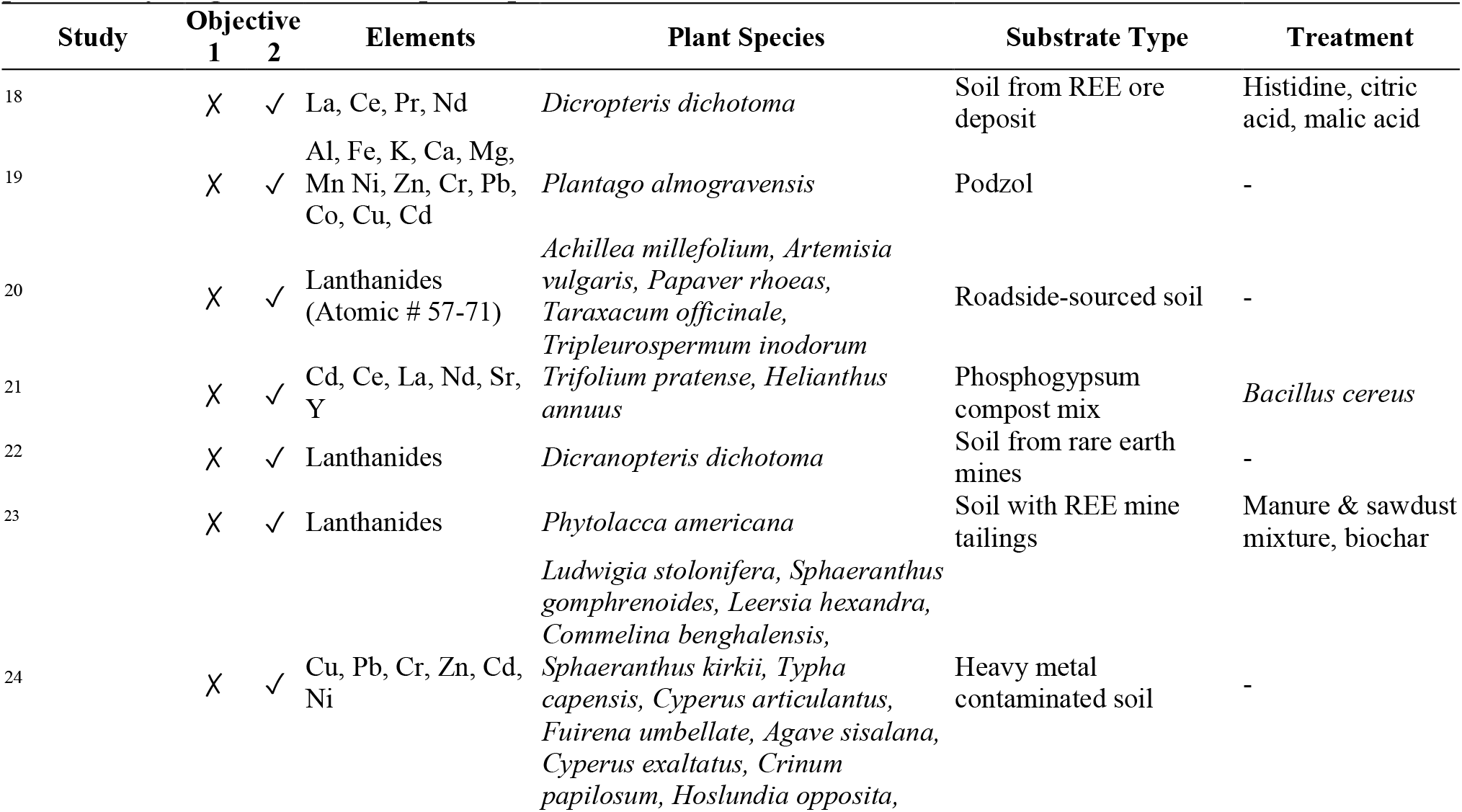
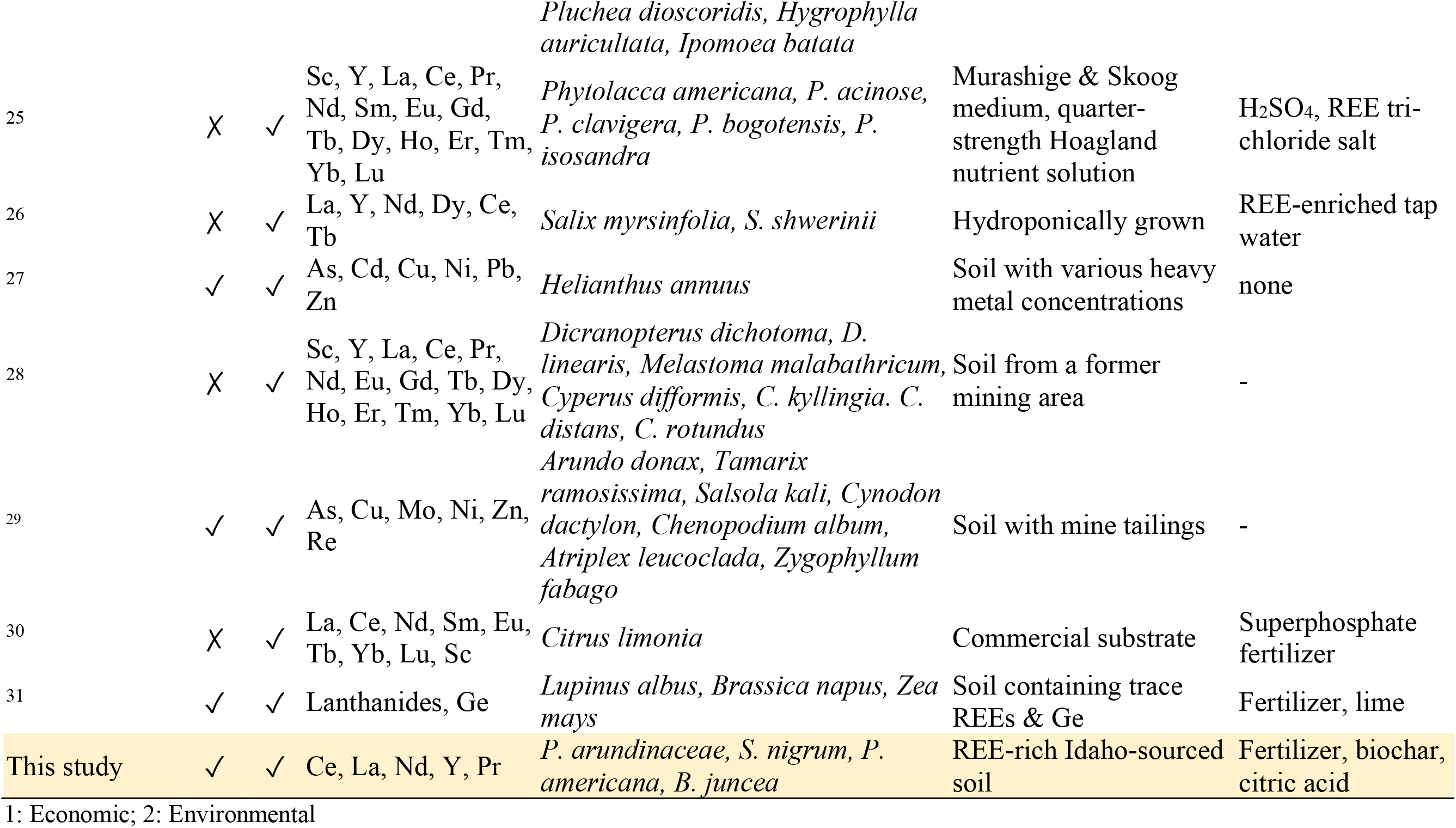
An overview of phytomining studies from the past two decades summarizing their research focus, particularly targeted elements, plant species, and treatments.

In this study, we focused on the hyperaccumulation ability of four plant species due to their high BF values displayed during our primary experimental studies and in previous literature (Supplementary **Table S2**). The selected plant species are *Phalaris arundinacea* (common name reed canary grass), *Solanum nigrum* (black nightshade), *Phytolacca americana* (pokeweed), and *Brassica juncea* (brown mustard) ^23,32–34^. To better understand how different environmental conditions impact hyperaccumulator performance, each plant species was subject to control conditions, soil treatment with fertilizer or biochar, and water treatment with citric acid to determine the best combination with REE-rich soil substrate. Biomass then underwent drying and pyrolysis to concentrate REEs for elemental analysis and assess the effectiveness of these steps.

## Results

### Growing success and bio-ore mass production

Out of the 18 individual pots planted per species, a pot was considered a success if enough biomass was produced to be harvested for pyrolysis and characterization (>0.1 g bio-ore). The overall success rates were as follows: 22% for *P. arundinacea*, 50% for *P. americana*, and 61% for *S. nigrum* and *B. juncea* (**Table 2**). Overall, 35 out of 72 pots were counted as successes. Multiple pots contained seeds that did not grow or failed to produce enough biomass for characterization. Fertilizer use was positively correlated with pot success rate (p<0.007) and biomass (p<0.005). Non-treated water was also associated with a higher success rate.

**Table 2.**
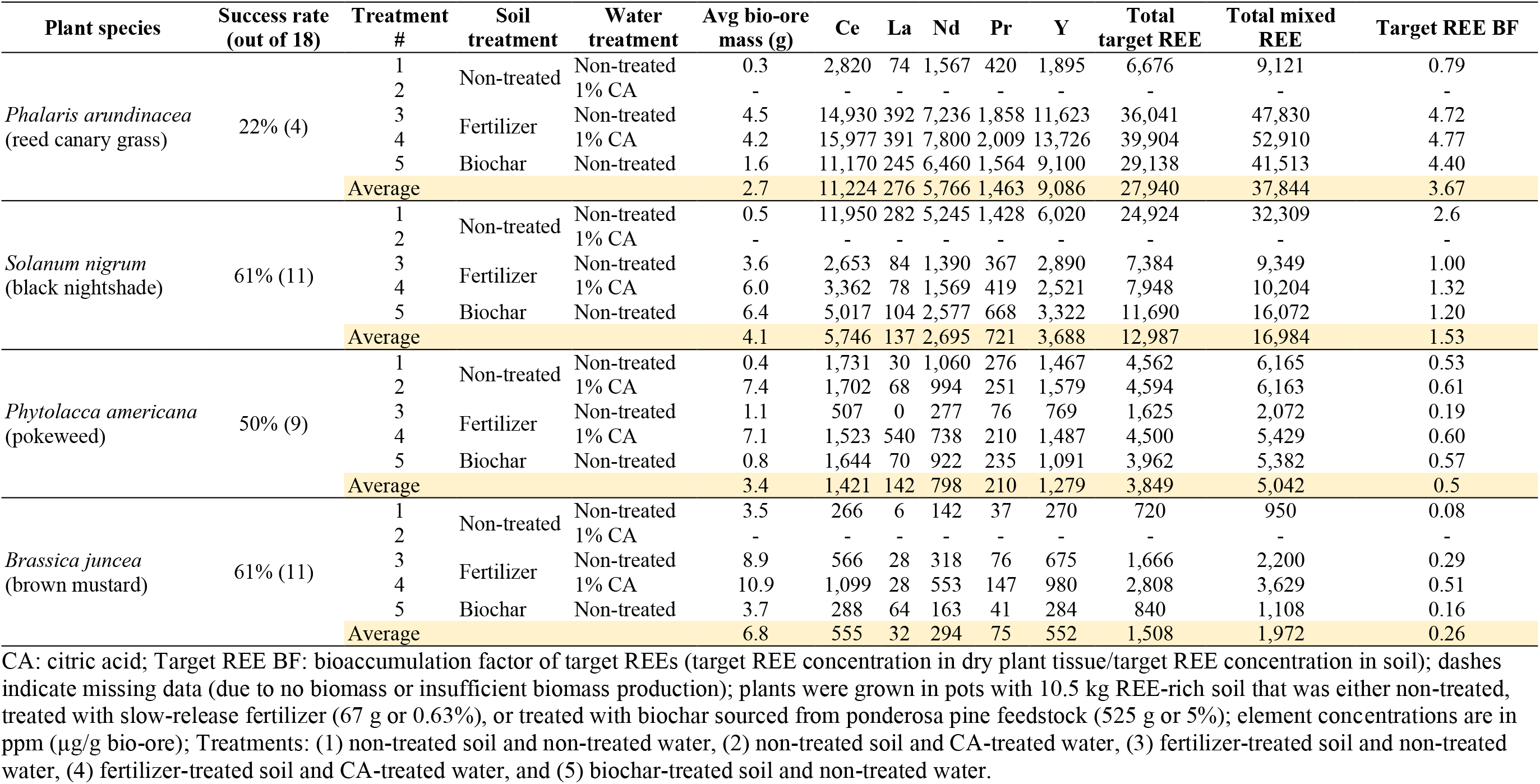
Summary of plant accumulation of REEs according to species and treatment.

During the initial growing period, individuals from all species except for *B. juncea* successfully grew from seeds into mature plants in the greenhouse (**Figure 1**). *B. juncea* individuals that had been planted as seeds directly into the 10.5 kg pots during the first round of planting did not germinate. It is assumed that the seeds did not receive sufficient warmth and light. To address this assumption, seeds were planted in growth trays containing the same soil type and stored in a warmer, brighter growth chamber before being transferred to 10.5 kg pots in the greenhouse, as detailed in the *Seed preparation* subsection in the *Methods* section. This vastly improved germination rate and may have contributed to the species’ considerable biomass production.

**Figure 1.**
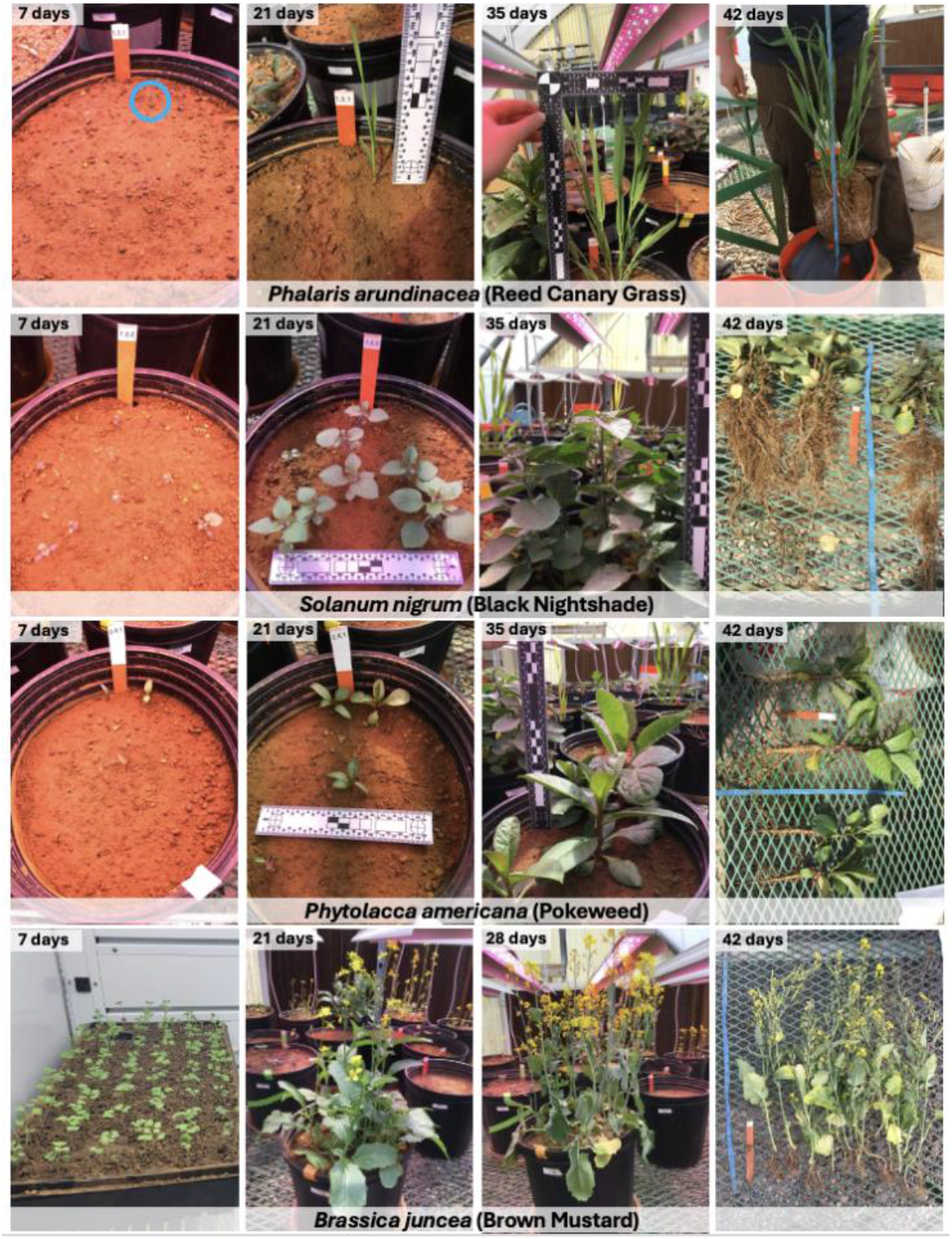
Growing stages of each of the four studied plant species from germination to harvest.

Treatment of soil with fertilizer or biochar increased bio-ore yield in grams (p < 0.001) (**Figure 2**). Supplementary **Table S3** presents the two-tailed generalized linear model results of bio-ore mass analysis. We found that soil treatment with fertilizer and water treatment with citric acid significantly affected bio-ore yield. Plant species did not significantly affect bio-ore mass production, except in the case of *B. juncea*. The REE concentration detected in the bio-ore did not considerably affect its production. Fertilizer-treated soil resulted in significantly higher bio-ore yields on average than the other soil treatments. Overall, *B. juncea* (brown mustard) produced the most bio-ore while *P. arundinacea* (reed canary grass) produced the least.

**Figure 2.**
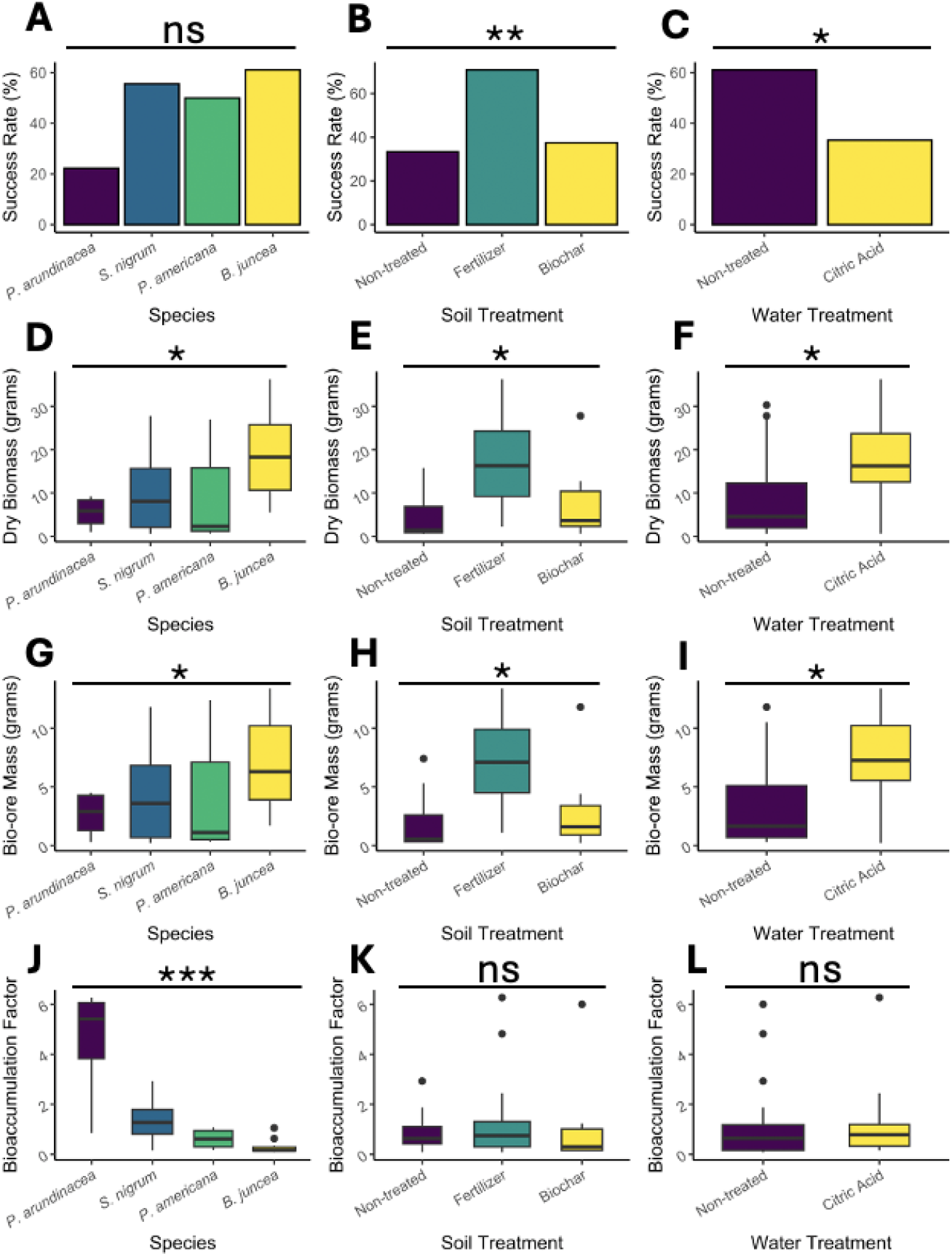
Boxplot summaries of the trends in plant success rate, dry biomass weight, bio-ore weight, and bioaccumulation factor. Boxplot width reflects the number of samples in that group (indicated in Table 3). Plant species are *Phalaris arundinacea, Solanum nigrum, Phytolacca americana*, and *Brassica juncea*. Soil treatments include no treatment, 0.63% fertilizer treatment, and 5% biochar treatment. Water treatments include no treatment and treatment with 1% citric acid. Plots show (A) plant success rate (%) according to species, (B) plant success rate according to soil treatment, (C) plant success rate according to water treatment, (D) dry biomass yield (in grams) according to species, (E) dry biomass yield according to soil treatment, (F) dry biomass yield according to water treatment, (G) bio-ore mass yield (in grams) according to species, (H) bio-ore mass yield according to soil treatment, (I) bio-ore mass yield according to water treatment, (J) bioaccumulation factor according to species, (K) bioaccumulation factor according to soil treatment, and (L) bioaccumulation factor according to water treatment. *, **, and *** indicate p<0.05, p<0.01, and p<0.001, respectively. “ns” indicates no significant relationship between factors (α = 0.05). Plots A-I show GLM test results. Plots J-L show ANOVA test results.

### Hyperaccumulation ability

Results indicate that *P. arundinacea* (reed canary grass) was the most effective hyperaccumulator of the four species tested, accumulating approximately 27,940 ppm (micrograms of REEs per gram of bio-ore) of mixed targeted REEs (Ce, La, Nd, Pr, Y) and exhibiting an average BF of 3.67. Supplementary **Table S4** provides plant accumulation of the targeted REEs, as well as the total accumulation of all present REEs. The second most effective hyperaccumulator was *S. nigrum* (black nightshade), accumulating approximately 12,987 ppm mixed targeted REEs, and displaying an average BF of 1.53. *P. americana* (BF of 0.5) and *B. juncea* (BF of 0.26) displayed weaker REE accumulation abilities.

**Table 3.** Averaged characteristics of collected soil from the Diamond Creek project site near Salmon, Idaho.

| Parameters | Values | Parameters | Values |
| --- | --- | --- | --- |
| pH | 7.6 | Calcium (meq/100g) | 12.4 |
| Sand % | 47.3 | Magnesium (meq/100g) | 3.4 |
| Silt % | 39.5 | Sulfate-S (ppm) | 3.0 |
| Clay % | 13.2 | Zinc (ppm) | 1.3 |
| Salts (mmhos/cm) | 0.6 | Iron (ppm) | 9.2 |
| Chlorides (ppm) | 7.0 | Manganese (ppm) | 1.5 |
| Sodium (meq/100g) | 0.2 | Copper (ppm) | 0.1 |
| CEC (meq/100g) | 16.3 | Boron (ppm) | 0.2 |
| Excess lime (%) | 3.2 | Cerium (ppm) | 2,889 |
| Organic matter (%) | 2.8 | Lanthanum (ppm) | 2,071 |
| Organic N (lb/Acre) | 55.0 | Neodymium (ppm) | 1,690 |
| Ammonium-N (ppm) | 2.6 | Praseodymium (ppm) | 435 |
| Nitrate-N (ppm) | 5.0 | Yttrium (ppm) | 808 |
| Phosphorous (ppm) | 5.0 | Total mixed target REEs (ppm) | 7,893 |
| Potassium (ppm) | 97.0 |  |  |

Soil and water treatments increased bio-ore production. However, these factors do not appear to play a significant role in BF. The ability to accumulate REEs is notably different among species. Supplementary **Table S5** provides the ANOVA result of BF analysis. Results from the ANOVA analysis performed indicated that species selection and the two-way interaction between species and soil treatment had significant effects on the uptake and storage of all target REEs except for La (Supplementary **Table S6**).

## Discussion

This study evaluated four plant species for their survival and ability to hyperaccumulate REEs (especially Ce, La, Nd, Pr, and Y) when grown in Idaho-sourced REE-rich soil. Plants were subjected to different soil (non-treated, fertilizer, and biochar) and water (non-treated and citric acid) treatments to additionally investigate the effects of differing conditions on bio-ore yield and bioaccumulation factor (BF). Fertilizer and biochar were selected as soil treatments due to evidence of their ability to promote plant tissue growth and increase hyperaccumulator yield via mobilization of target elements ^32,35^. Citric acid was chosen as a water treatment based on previous studies that show that small doses enhance REE uptake via the root system, aid in mobilization for transfer from the roots to aboveground tissues, decrease plant stress, and boost biomass production ^10,36,37^.

Results show that plant species had a significant effect on BF, with *Phalaris arundinacea* displaying the highest value of all tested species. *Brassica juncea* produced the most bio-ore, which may have been due to the different germination conditions where seeds sprouted in a growth chamber rather than directly in the greenhouse pots. The insignificant impact of soil and water treatments on BF conflicts with the results of Turra et al., who found that fertilizer application and decrease of pH (in this study via citric acid application) result in higher plant tissue REE concentration ^38^. Regarding bio-ore mass production, soil and water treatments appeared to play a more significant role than species type in our study. To maximize the effectiveness and efficiency of a phytomining operation, it is important to find a balance between hyperaccumulation ability and bio-ore (and biomass) yield. A high BF is essential for extracting target REEs, but bio-ore yield must also be considerable as it is what ultimately determines how much of the resource will be recovered from the soil. Since REE hyperaccumulators are not well described as compared to those of other metals (e.g., plants are considered nickel hyperaccumulators when they concentrate at least 1,000 ppm in their dried tissues), this study acts as a valuable contribution to the limited existing body of knowledge ^39^. The research into the impact of different treatments additionally aids in the understanding of how to increase bio-ore production during the growing process.

A major challenge presented in this experimental study was the failure of multiple plant individuals to germinate or to gain biomass after sprouting. Previous research has shown that an overabundance of metals in the soil can cause a decline in root development, interference with mitosis, and a decrease in biomass production, among other problems ^40^. Soil pH that maintains plant health tends to fall between 6.2 and 6.8, and the REE-rich soil exhibits a far more basic average pH of 7.6, which could have impacted plant health, reducing biomass yield and survival rate ^41^. Fertilizer significantly increased pot success rate and bio-ore mass production. These factors pose the question of what amendments (e.g., fertilizer, biochar, or citric acid) should be made to this substrate to make it more hospitable for future use.

The proposed low-emission pathway (i.e., phytomining, drying, and pyrolysis) is an environmentally sustainable approach for REE extraction from REE-rich surface soil, especially when using the plant species *P. arundinacea*. To date, few studies have explored hyperaccumulator effectiveness on REE-rich substrate, and none have used Idaho-sourced soil. Our research serves as a foundation for future investigation and improvement of the proposed extraction pathway on REE-rich surface soils and similar substrates. However, the narrow scope of this study warrants further investigation into various applications of hyperaccumulator species and performance-enhancing treatments.

## Methods

### Soil preparation

Soil was collected from the Diamond Creek project site (located near Salmon, Idaho) in the summer of 2022 (**Figure 3**). Rocks were removed from the soil with a #5 mesh (0.157 inches or 4 millimeters) and triturated into finer grains using a mixer and loose metal ball bearings. The resulting material was then homogenized back into the fine soil using the same mixer.

**Figure 3.**
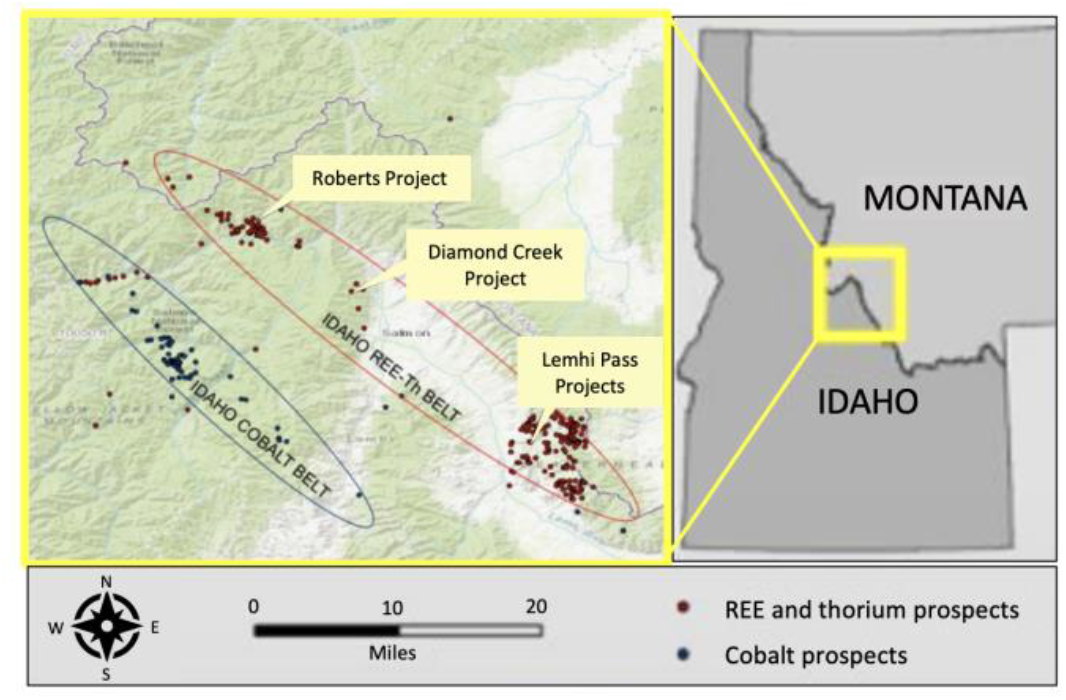
Map of REE deposits and other valuable mineral prospects in central Idaho (Courtesy of Idaho Strategic Resources, Inc.).

**Table 3** provides the soil characteristics and elemental results (analytical methods detailed in Supplementary **Table S7**). Planting pots were filled with approximately 10.5 kg of soil and characterized using an X-ray fluorescence (XRF) spectrometer to determine and record the average composition of targeted REEs, i.e., cerium (Ce), lanthanum (La), neodymium (Nd), praseodymium (Pr), and yttrium (Y).

The soil in each pot was either treated with 67 grams of fertilizer, 525 grams of biochar, or left non-treated (**Table 2**).

### Seed preparation

Packaged, commercially available seeds of the four chosen plant species *Phalaris arundinacea* (reed canary grass), *Solanum nigrum* (black nightshade), *Phytolacca americana* (pokeweed), and *Brassica juncea* (brown mustard) were obtained and used in this study. Prior to sowing, seeds of *P. americana* were subjected to cold-stratification by storing in a humid environment at 5 °C for 3-4 weeks, followed by incubation in a growth chamber at 26.5 °C until germinated (approximately seven days). Seeds of *B. juncea* were similarly incubated in a growth chamber at 26.5 °C for seven days until germination. After preparing *P. americana* and *B. juncea* seeds, all four species were sown in the prepared pots. Each pot received 11 seeds or seedlings of the singular species randomly assigned to it.

### Experimental design, growing, and treatments

The experiment was established in a greenhouse as a randomized complete block design with three blocks (replications) and three factors: plant species (four levels: *P. arundinacea, S. nigrum, P. americana*, and *B. juncea*), soil treatment (three levels: non-treated, fertilizer-treated, and biochar-treated), and water treatment (two levels: non treated and citric acid-treated). Overall, 72 pots were prepared for planting with 18 assigned to each species. Each unique combination of species, soil treatment, and water treatment was replicated three times. Seeds were planted in pots containing REE-rich soil from the Diamond Creek site that was non-treated or treated with either Osmocote® (a slow-release fertilizer with a nitrogen-phosphorous-potassium ratio of 14:14:14) or biochar (nutrient-rich pyrolyzed material from pinewood). The biochar used in this study was sourced from ponderosa pine (*Pinus ponderosa*) feedstock. Each pot was subjected to either non-treated tap water or tap water treated with 1% citric acid. Laboratory results indicated the water had a pH of 7.6 and hardness of 14.76 grains per gallon. Pots were bottom watered (water poured into a tray beneath the pot) with 300 mL and top watered (water poured on top of the soil in the pot) with 300 mL of the assigned water solution three times per week with at least one full day in between waterings. Plants were grown in a greenhouse that was consistently kept between 18 °C and 32 °C. Grow lights were used in conjunction with natural light to maintain a minimum of 308 µmol/m^2^/s for 16 hours a day, followed by 8 hours of darkness.

### Plant harvesting

After six weeks of growth, plants that survived were harvested. Individual plants were gently removed from pots by hand to recover as much underground root biomass as possible. Specimens were washed with tap water to remove all excess soil. Roots were separated from the aerial parts and gently agitated in an ultrasonic cleaner to ensure the removal of as many soil particles as possible. Following washing, the total biomass collected from each pot was weighed and recorded. Tissues were left to dry for 24 hours in a 95 °C oven and then finely ground using a mortar and pestle. Plant biomass was weighed once dried to estimate initial moisture content.

### Pyrolysis and elemental analysis

Dry biomass was placed into a small reactor and pyrolyzed at 400 °C for 30 minutes while nitrogen gas was pumped into the reaction chamber at 1.5-3.0 LPM to displace any oxygen gas. Three distinct products were formed in this process: pyrolysis oil, gas, and REE-rich solid (bio-ore). Pyrolysis oil was weighed, but not collected or analyzed.

Pyrolysis gas was released without being collected. Bio-ore samples were weighed and prepared using the United States EPA 3050B acid digestion method to concentrate REEs into a liquid suspension for detection ^42^. The 5 mL resulting product was run through a 45-micrometer filter via vacuum filtration to remove solids and diluted with ultrapure water to 100 mL. The diluted product then underwent inductively coupled plasma mass spectrometry (ICP-MS) to detect concentrations of REEs that the plants in each pot took up collectively. Depending on the bio-ore amount, up to three digestions were made for each sample, and each digestion was analyzed twice to maximize measurement repetition.

### Calculations and statistical analysis

Once REE composition in µg/L was obtained via ICP-MS characterization, the results were converted to ppm (µg/g). Target REE bioaccumulation factor (BF) was calculated for each sample by dividing the total ppm of targeted REEs in the dried plant tissue by the total ppm of targeted REEs in the Diamond Creek site soil (**Eq. 1**). BF represents the efficiency of a species and treatment combination in extracting and storing the target REEs from the soil and retaining these materials through processing via pyrolysis, with higher numbers indicating higher extraction and retention ability.

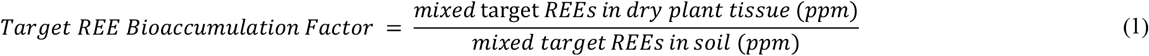

Statistical analysis was performed, using R (an open-source programming language) ^43^. BF was normalized and modeled as a normally distributed variable, while bio-ore mass was modeled as a gamma-distributed variable. A two-tailed analysis of variance (ANOVA) was performed to analyze the significance of the key parameters (i.e., plant species, soil treatment, and water treatment) on BF. In the case of bio-ore mass analysis, a two-tailed generalized linear model with a log link function was used. The significance level was drawn at α = 0.05 for both approaches.

## Acknowledgements

The authors wish to acknowledge Idaho Department of Commerce (IGEM-Commerce Grant #5358) for funding this project and Idaho Strategic Resources Inc. (IDR) for their resources and support. We would also like to thank the University of Idaho, Idaho Falls Research and Extension Center for their resources, support, and use of their greenhouse.

## Author Contributions

**Kathryn Richardson:** Methodology, Investigation, Formal Analysis, Writing – Original Draft, Visualization. **Amin Mirkouei:** Conceptualization, Methodology, Writing – Review and Editing, Supervision, Project Administration, Funding Acquisition. **Kasia Duellman:** Resources, Writing – Review and Editing, Supervision. **Anthony Aylward:** Visualization, Writing – Review and Editing. **David Zirker:** Conceptualization, Methodology, Investigation, Writing – Review and Editing. **Eliezer Schwarz:** Writing – Review and Editing. **Ying Sun:** Writing – Review and Editing.

## Declaration of Competing Interest

The authors declare that they have no known competing financial interests or personal relationships that could have appeared to influence the work reported in this paper.

## Data Availability

The authors declare that data used to reach the findings of this study are provided in this manuscript, Supplementary Information, and GitHub (https://github.com/RSMLResearchGroup/Phytomining). Additional data are available upon request.

## Code Availability

The codes are available on GitHub (https://github.com/RSMLResearchGroup/Phytomining).

## Supplementary Information

**Table S1.** Average concentration of each REE in soil from Dimond Creek project site near the town of Salmon in Idaho, USA.

| Element | Ce | Dy | Er | Eu | Gd | Ho | La | Lu | Nd | Pr | Sc | Sm | Tb | Tm | Y | Yb | Total |
| --- | --- | --- | --- | --- | --- | --- | --- | --- | --- | --- | --- | --- | --- | --- | --- | --- | --- |
| Value (ppm)* | 4,338 | 309 | 101 | 138 | 431 | 47 | 1,865 | 6 | 2,052 | 505 | 12 | 447 | 60 | 11 | 1,226 | 53 | 11,601 |

\* Data obtained via X-ray fluorescence (XRF) spectrometer. Element concentrations are presented in $\mu\text{g/g}$ (ppm).

**Table S2.** Summary of primary experimental studies on seven plant species. Seeds were planted directly into pots filled with 2.5 kg Lucky Gem soil and kept moist until germination. Germinated plants were top and bottom watered three times per week with a total of 1,350 mL non-treated tap water and allowed to grow for 20 weeks until harvest. Biomass was harvested, washed with tap water, pyrolyzed at 400 °C for 30 minutes, acid digested, and characterized via ICP-MS.

| Species Used | $\mu\text{g/g REE}$ | | | Bioaccumulation Factor | | |
| --- | --- | --- | --- | --- | --- | --- |
|  | La | Ce | Nd | La | Ce | Nd |
| Brassica juncea (brown mustard) | 180 | 430 | 290 | 0.32 | 0.39 | 0.45 |
| Helianthus annuus (honeycomb sunflower) | 30 | 62 | 34 | 0.0014 | 0.0017 | 0.00093 |
| Helianthus annuus (hybrid 894 sunflower) | 0.86 | 2 | 1.3 | 0.0015 | 0.0018 | 0.0020 |
| Medicago sativa (alfalfa) | 3.5 | 6.8 | 4.5 | 0.0062 | 0.0061 | 0.0070 |
| Phytolacca americana (pokeweed) | 30 | 62 | 34 | 0.053 | 0.056 | 0.053 |
| Solanum nigrum (black nightshade) | 130 | 280 | 180 | 0.23 | 0.25 | 0.28 |
| Verbascum thapsus (common mullein) | 43 | 91 | 46 | 0.076 | 0.082 | 0.071 |
| Soil | 560 | 1100 | 640 | - |  |  |

**Table S3.** P-value results for the two-tailed generalized linear models run to analyze plant success rate, dry biomass, and bio-ore mass.

| Factor | Success rate | Dry mass | Bio-ore mass |
| --- | --- | --- | --- |
| Intercept | 0.96629 | 0.1458 | 0.8403 |
| Fertilizer-treated soil | 0.73725 | 0.1331 | 0.0491 |
| Biochar-treated soil | 0.01047 | 0.0367 | 0.1913 |
| Citric acid-treated water | 0.70474 | 1.049 | 0.0434 |
| *Phalaris arundinacea* | 0.44908 | 0.0121 | -0.114 |
| *Solanum nigrum* | 0.000618 | 0.0376 | 0.842 |
| *Brassica juncea* | 0.73725 | 0.1331 | 0.0416 |

**Table S4.**
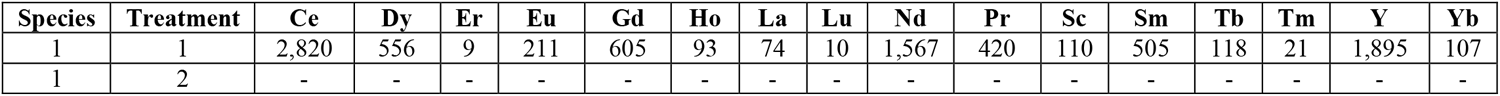
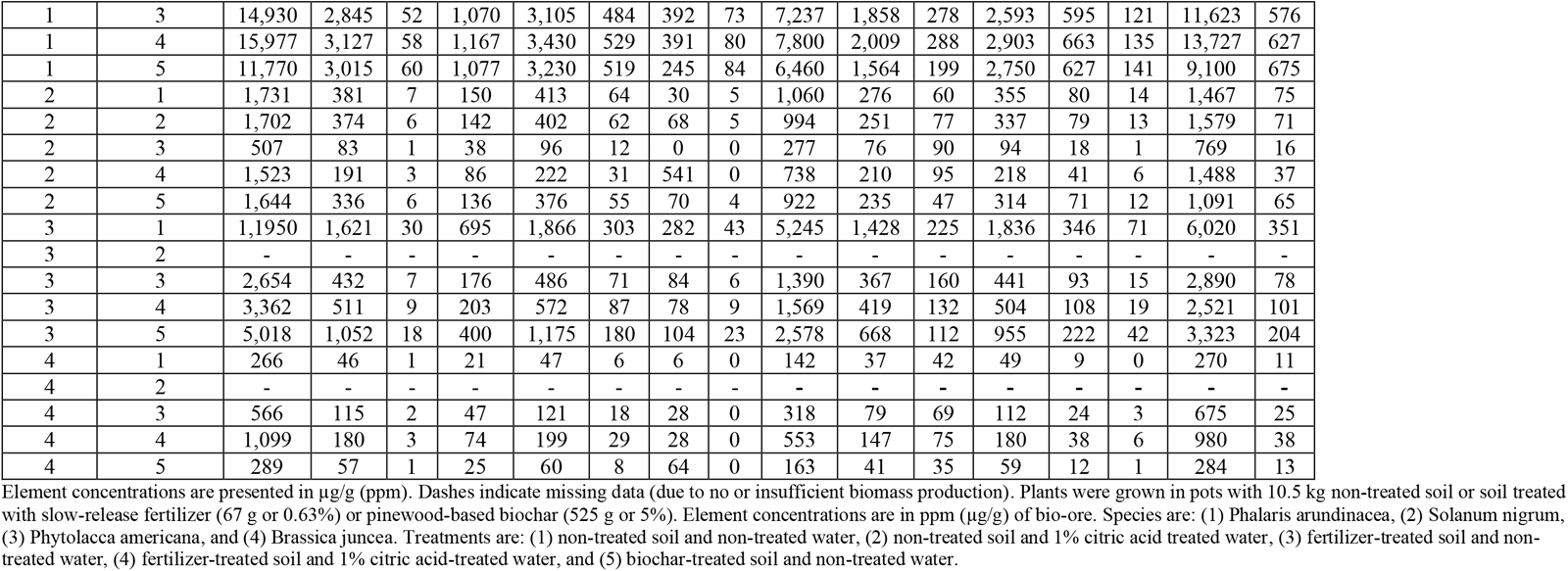
Plant accumulation of all REEs for various plant species and treatments. Element concentrations are presented in µg/g (ppm). Dashes indicate missing data (due to no or insufficient biomass production). Plants were grown in pots with 10.5 kg non-treated soil or soil treated with slow-release fertilizer (67 g or 0.63%) or pinewood-based biochar (525 g or 5%). Element concentrations are in ppm (µg/g) of bio-ore. Species are: (1) Phalaris arundinacea, (2) Solanum nigrum, (3) Phytolacca americana, and (4) Brassica juncea. Treatments are: (1) non-treated soil and non-treated water, (2) non-treated soil and 1% citric acid treated water, (3) fertilizer-treated soil and non-treated water, (4) fertilizer-treated soil and 1% citric acid-treated water, and (5) biochar-treated soil and non-treated water.

**Table S5.** ANOVA results of bioaccumulation factor analysis.

| Parameters | P-value |
| --- | --- |
| Species (main effect) | 1.04e-09 |
| Soil treatment (main effect) | 0.180 |
| Water treatment (main effect) | 0.448 |
| Species: Soil treatment (2-way interaction) | 1.34e-05 |
| Species: Water treatment (2-way interaction) | 0.328 |
| Soil treatment: Water treatment (2-way interaction) | 0.165 |

**Table S6.** ANOVA results of ICP-MS REE ppm values.

| Factor | P-value |  |  |  |  |  |
| --- | --- | --- | --- | --- | --- | --- |
|  | Ce | La | Nd | Pr | Y | Mixed REEs |
| Species (main effect) | 1.09e-06 | 0.334 | 7.98e-08 | 2.28e-07 | 1.27e-10 | 4.59e-08 |
| Soil (main effect) | 0.538541 | 0.712 | 0.658876 | 0.549465 | 0.110 | 0.606 |
| Water (main effect) | 0.750162 | 0.663 | 0.644551 | 0.683979 | 0.638 | 0.714 |
| Species: Soil treatment (2-way interaction) | 0.000373 | 0.561 | 0.000126 | 0.000223 | 1.16e-06 | 0.644e-05 |
| Species: Water treatment (2-way interaction) | 0.854462 | 0.735 | 0.701355 | 0.729351 | 0.110 | 0.612 |
| Soil treatment: Water treatment (2-way interaction) | 0.316128 | 0.614 | 0.212972 | 0.243298 | 0.281 | 0.263 |

**Table S7.** Analytical methods used by Stukenholtz Laboratory (Twin Falls, ID) to determine relevant soil and water characteristics.

| Characteristic | Analytical Method(s) |
| --- | --- |
| Soil pH | 1:1 (soil:water), method S-2.20* |
| Soil EC | Soil water 1:1, electrical conductivity 25 °C, method S-2.30* |
| Soil excess (free) lime | Sulfuric acid dissolution, NaOH back titration, modified method 955.01 Neutralizing Value for Liming Materials** |
| Soil organic matter | Gravimetric mass loss on ignition, 2 hrs 360 °C, method S-9.20* |
| Soil ammonium-N | 1:5 extraction KCl colorimetric determination, method S-3.50* |
| Soil nitrate-N | 1:5 extraction KCl, Cd-reduction, method S-3.10* |
| Soil phosphate-P<sub>Olsen</sub> | Olsen, 1:20 extraction, colorimetric determination 880 nm method S-4.10* |
| Soil phosphate-P<sub>Bray</sub> | Bray, 1:20 5 min extraction, colorimetric determination 880 nm method S-4.20* |
| Soil potassium, sodium, calcium, magnesium | Ammonium acetate 1:10 extraction, ICP determination, method S-5.10* |
| Chloride, sulfide, boron | AA-NH<sub>4</sub>F Kewlona extraction, ICP determination, method S-4.60* |
| Zinc, iron, manganese, copper | DTPA extraction, method S-6.10* |
| SAR (sodium absorption ratio) | SAR saturated paste extraction of Ca, Mg, Na, method S-1.60* |
| Water pH | Method W-1.10* |
| Water hardness | Method W-1.60* |

\* Method detailed in Soil, Plant, and Water Reference Methods for the Western Region <sup>44</sup>
\*\* Method detailed in Official Methods of Analysis of AOAC International <sup>45</sup>

## Notes

https://github.com/RSMLResearchGroup/Phytomining

### Competing Interest Statement

The authors have declared no competing interest.

